# Matching or genetic engineering of HLA Class I and II facilitates successful allogeneic ‘off-the-shelf’ regulatory T cell therapy

**DOI:** 10.1101/2023.08.06.551956

**Authors:** Oliver McCallion, Weijie Du, Viktor Glaser, Kate Milward, Clemens Franke, Jonas Kath, Mikhail Valkov, Mingxing Yang, Annette Künkele, Julia K. Polansky, Michael Schmueck-Henneresse, Hans-Dieter Volk, Petra Reinke, Dimitrios L. Wagner, Joanna Hester, Fadi Issa

## Abstract

The potential to harness regulatory T cells (Tregs) for the treatment of autoimmune diseases and transplant rejection has been restricted by several barriers: donor variability, manufacturing complications, and time-consuming expansion processes. These issues further complicate the use of autologous Tregs during acute disease phases or when Tregs are low in number or dysfunctional. Here we explore the potential of ‘off-the-shelf’ allogeneic Tregs, from healthy donors or universal sources, to provide a more practical solution. We discover that the efficacy of these cells is undermined by the recipient’s immune response, and that that rigorous matching of HLA classes I and II overcomes this barrier. Importantly, genetically manipulating HLA expression enables the use of unmatched allogeneic Tregs with *in vivo* efficacy. Our findings underscore the transformative potential of HLA-engineered Tregs, offering a novel, ready-to-use therapeutic avenue for treating a wide array of inflammatory diseases.

**One-Sentence Summary:** Matching or engineering of HLA-I and HLA-II facilitates allogeneic ‘off-the-shelf’ regulatory T cells for immunoregulation.

## Main Text

Regulatory T cells (Tregs) are potent immunosuppressive lymphocytes defined by expression of the FOXP3 transcription factor. Significant progress towards harnessing the therapeutic potential of Tregs to dampen pathological immune activity has been achieved with several recent successes in autoimmunity, graft-versus-host disease (GvHD), and transplant rejection (*1-8*).

To date, clinical trials have predominantly investigated adoptive cell transfer approaches. Here, cell therapy recipients donate blood or tissue from which Tregs are isolated, purified, and expanded *ex vivo* to create a bespoke infusion that is returned to the patient (*9*). There are several challenges to this autologous approach. Firstly, individuals harbor variable numbers of Tregs, particularly at the extremes of age, which potentially limits pre-expansion starting numbers and may restrict patients for whom autologous cell therapy is possible. Equally, the underlying pathology for which Treg therapy is required may be associated with impaired Treg function (*10-13*). On the manufacturing side, Treg expansion takes several weeks to complete and may fail from biological or technical complications including poor Treg expansion or contamination with non-Treg cells (*14*). These manufacturing constraints preclude the acute use of autologous regulatory cell therapies, for example in transplant recipients of deceased donor organs or during rejection episodes or autoimmune disease flare-ups. Finally, named-patient cell products require significant time, resources, and expertise to produce, reflected in significant manufacturing cost which will likely limit their broad availability for patients (*15*).

For these reasons, the prospect of producing pre-manufactured cell therapy products from healthy human donors or a universal cell source, such as induced pluripotent stem cells (iPSCs), is appealing (*16, 17*). An ‘off-the-shelf’ solution would facilitate selection and scaled production of maximally suppressive Tregs and would equally offer increased flexibility through providing a product available for acute, unscheduled administration, potentially enabling investigation of regulatory cell therapy for a broader range of pathologies (*16*). Furthermore, the scaled manufacture of a cell product provides the ideal platform for engineering and banking of next-generation therapeutic products. However, as with any allogeneic cell therapy, it is likely that the recipient immune system will reject infused cells, as already shown for allogeneic effector and CAR T cell therapies (*18-20*). It is not fully understood yet whether this is also the case for unmatched Treg because of their inherent immunosuppressive capacity (*21, 22*).

In this study we develop methodologies that permit the *in vivo* survival and function of allogeneic Tregs. *In vitro*, allogeneic Tregs demonstrate similar suppressive capabilities to autologous Tregs, highlighting the difficulties interpreting *in vitro* functional assays. *In vivo*, killing by host CD8^+^ T cells dramatically reduces the efficacy of allogeneic Tregs, even when infused at relatively high numbers. This killing is circumvented by either HLA-matching of Tregs to the host, or through the CRISPR-Cas9 silencing of HLA-class I and II in the infused Treg cell therapy product. To further refine this approach, the replacement of polymorphic HLA-class I with a non-polymorphic HLA-E-*B2M* fusion gene (*23, 24*) restricts Treg elimination by NK cells. Together, these techniques define a set of successful strategies for effective ‘off-the-shelf’ Treg cellular therapy.

## Results

### Allogeneic Tregs do not lose in vitro suppressive capacity but are markedly less effective than autologous Tregs in vivo

To compare the ability of Tregs to suppress either autologous or allogeneic responders, we assessed their function using standard *in vitro* suppression assays (*25*) in which suppression of proliferation of responder PBMCs is assessed in the presence or absence of Tregs. Autologous and allogeneic Tregs suppressed responder cell proliferation equally (**Fig. 1A, B, Suppl. Fig. S1**). There were no significant differences in expression of the activation marker CD25 between responder cells suppressed by autologous compared to allogeneic Tregs (**Fig. 1C, 1D**). These results demonstrate that Treg suppress allogeneic and autologous effector cells equally well *in vitro*.

**Fig. 1.**
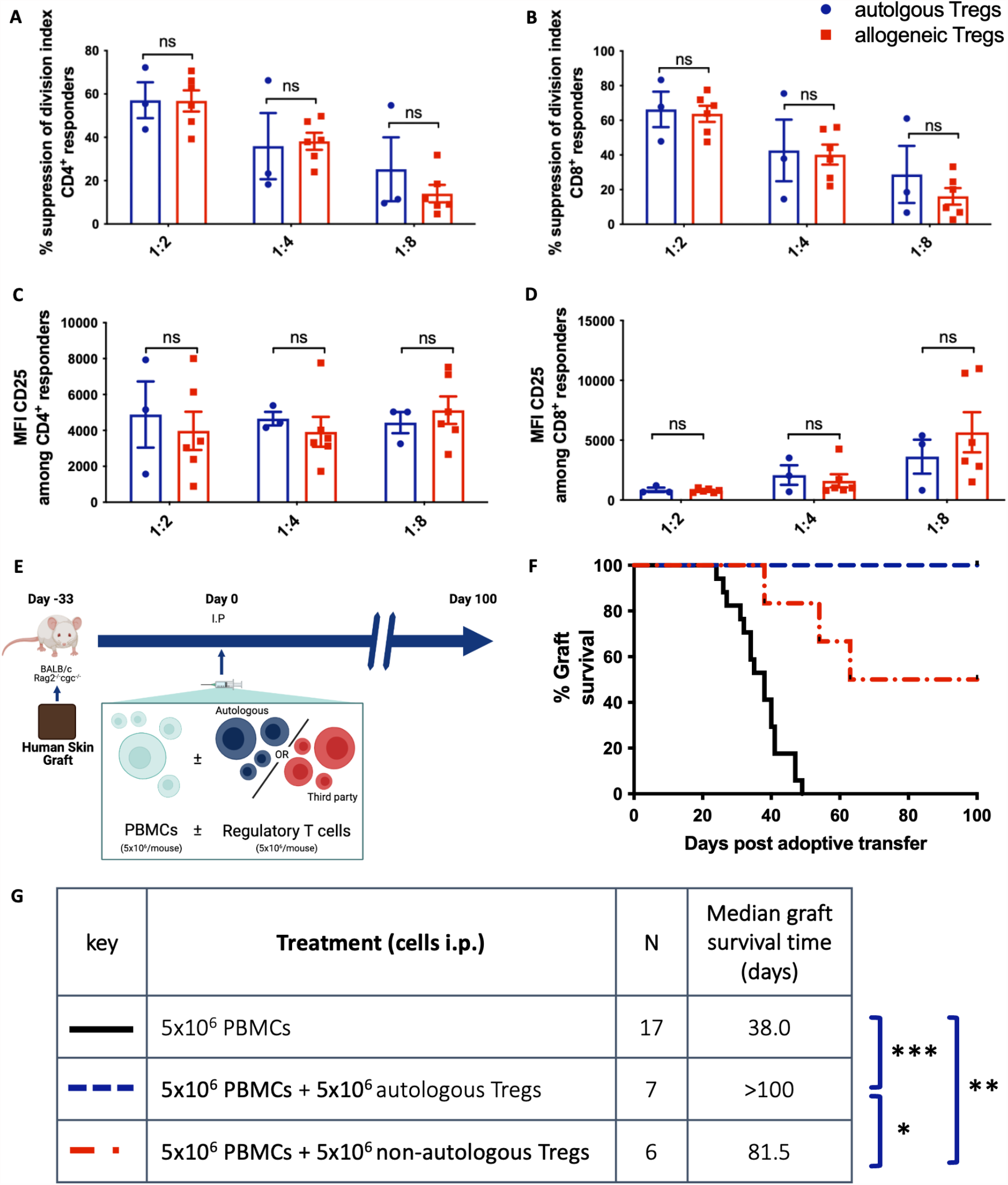
Autologous and allogeneic Tregs suppress effector proliferation *in vitro* and protect human skin graft rejection *in vivo*. CSFE-stained human PBMCs (1×10^5^) were stimulated with αCD3/αCD28-coated beads (2×10^4^) with and without either autologous or allogeneic Tregs (at 1:2, 1:4 and 1:8 Treg:PBMC ratios). CSFE dilution after 72hrs incubation was measured by flow cytometry and a division index was calculated. (**A**) The percentage suppression of CD4^+^ proliferation p=0.2495 (p>0.05 for all pairs). (**B**) The percentage suppression of CD8^+^ proliferation p=0.5897; (p>0.05 for all pairs). Both are calculated relative to stimulated PBMCs cultured in the absence of Tregs. (**C**) Median fluorescence intensity of CD25 amongst CD4^+^ responder cells p=0.7178 (p>0.05 for all pairs). **(D)** Median fluorescence intensity of CD25 amongst CD8^+^ responder cells p=0.5977 (p>0.05 for all pairs). Data are plotted as mean ± SEM of 3 autologous and 6 allogeneic responder:Treg donor combinations. All assays were performed in triplicate. Statistical significance for autologous versus allogeneic Tregs, across all Treg:responder ratios, was assessed using two-way repeated measures ANOVA with Bonferroni tests for pairwise comparisons. (**E**) Immunodeficient BALB/c Rag2^-/-^ cγc^-/-^ mice grafted with human skin received intraperitoneal human PBMCs (5×10^6^) alone (n=17), with Tregs autologous to the PBMC donor (5×10^6^, n=7) or Tregs allogeneic to the PBMC donor (5×10^6^, n=6). Grafts were monitored for macroscopic signs of rejection over the following 100 days. (**F**) Percentage of grafts surviving is plotted over time post-adoptive transfer of cells. (**G**) Sample size and median survival time for each treatment group is tabulated. Statistical significance was assessed using Mantel-Cox log rank tests: *p=0.0387; **p=0.0006; ***p<0.0001.

To determine the capacity for allogeneic Tregs to suppress the *in vivo* alloresponse, we utilized a mouse model of human skin transplant rejection (*25*) (**Fig. 1E**). Transplant survival was significantly prolonged by Treg treatment, although allogeneic Tregs demonstrated reduced efficacy in comparison to autologous cells (median graft survival time, MST, 81.5 vs >100 days) (**Fig. 1F, G**). This contrasted with *in vitro* suppression data, in which no differences were identified between the two Treg populations.

### Allogeneic Tregs are subject to profound CD8-mediated depletion in vivo

To investigate the mechanisms underlying this reduced *in vivo* efficacy, we subjected Tregs to both *in vitro* and *in vivo* cell survival assays. First, autologous or allogeneic Tregs were cultured with freshly isolated PBMCs. Treg survival was calculated as a proportion of the initial seeding density. In this *in vitro* assay, no significant differences in Treg survival were identified (**Fig. 2A**). For the *in vivo* assay, human PBMCs with or without CFSE-labelled Tregs were injected intra-peritoneally into immunodeficient mice and recovered after 7 days by peritoneal lavage (**Fig. 2B**). Here, there was a significantly higher recovery of autologous compared with allogeneic Tregs (**Fig. 2C**), suggesting reduced *in vivo* survival of Tregs in an allogeneic host. To determine the host cells responsible for allogeneic Treg loss, we compared Treg survival in mice receiving total PBMCs or PBMCs depleted of either CD8^+^ or CD56^+^ cells (**Suppl. Fig. S2**). Depletion of CD8^+^ T cells effectively restored the number and proliferative capacity of allogeneic Tregs recovered to a level comparable to autologous Tregs (**Fig. 2D-F**). Depletion of CD56^+^ cells did not have a similar impact, suggesting that the impaired survival and function of allogeneic unmodified Treg in this model is predominantly driven by CD8^+^ cells.

**Fig. 2.**
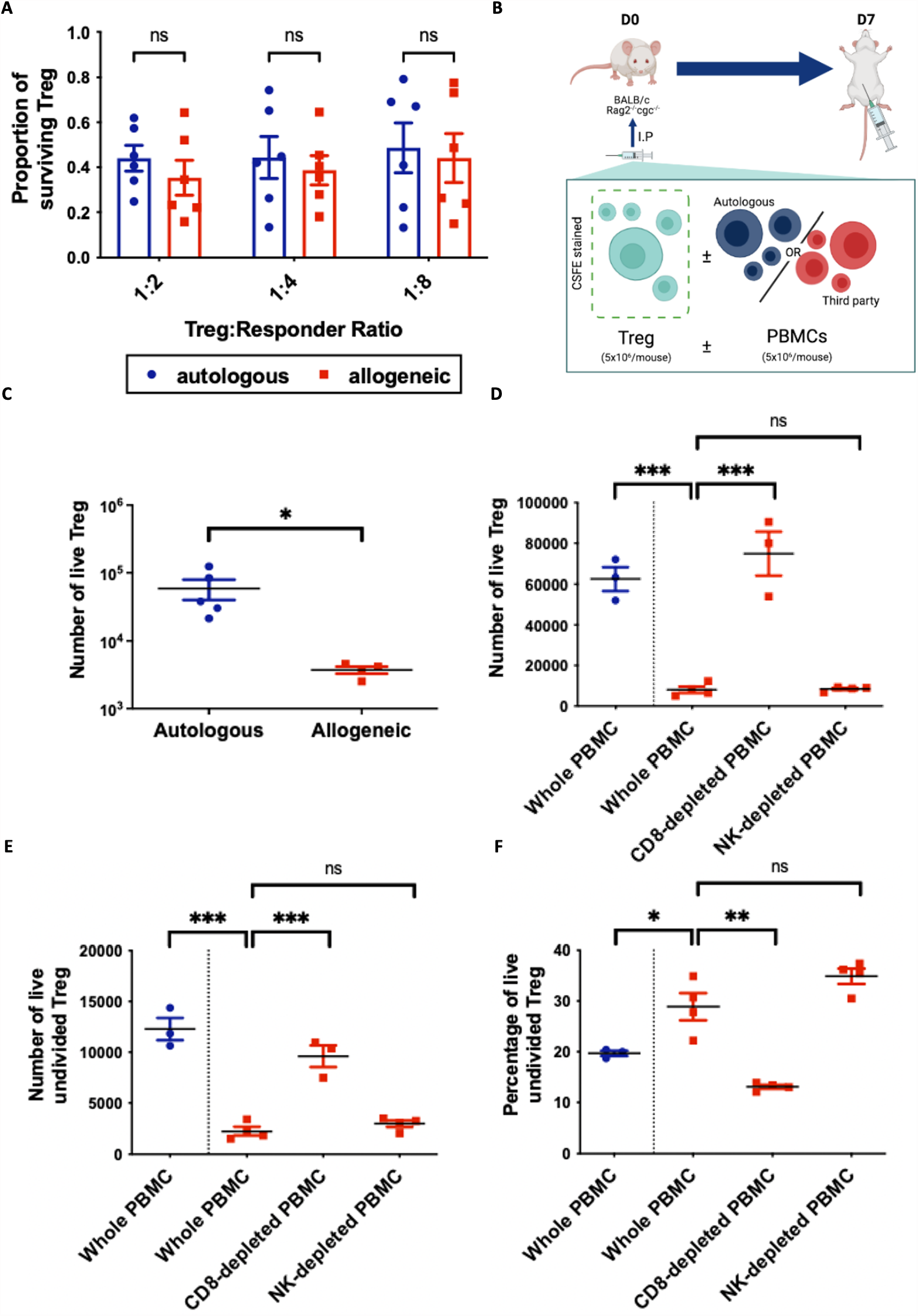
Autologous and allogeneic Treg survival is comparable *in vitro.* Allogeneic Treg survival is impaired *in vivo* which is reversed by depletion of host CD8^+^ cells. (**A**) Human *ex vivo*-expanded Tregs were cultured with VPD-stained autologous or allogeneic PBMCs (1×10^5^) in the presence of αCD3/αCD28-coated beads at Treg:PBMC ratios of 1:2, 1:4 and 1:8. After 3 days, the number of live Tregs in each well was enumerated by flow cytometry. Treg numbers are plotted as a fraction of the number of Tregs added on day 0. Data are presented as mean +/-SEM for 6 Treg donors. Statistical analyses, comparing Tregs cultured with autologous versus allogeneic PBMCs, were conducted using a two-way repeated measures ANOVA (across all Treg: PBMC ratios), with Bonferroni post-tests for Treg: PBMC ratio: p=0.9266 (two-way ANOVA); p>0.05 for each pair (Bonferroni). (**B**) Immunodeficient BALB/c Rag2^-/-^ cγc^-/-^ mice received intraperitoneal CFSE-stained human Tregs (5×10^6^) with 5×10^6^ autologous (n=5) or allogeneic (n=4) PBMCs. 7 days after injection, cells were recovered from the peritoneal cavity by lavage. CD4^+^CFSE^+^ Tregs in the effluent were enumerated by flow cytometry. (**C**) Cell numbers are plotted for each mouse individually, with mean ± SD for each treatment group. *p=0.0159, calculated using a Mann Whitney U test. (**D-F**) Immunodeficient BALB/c Rag2^-/-^ cγc^-/-^ mice received intraperitoneal CFSE-stained human Tregs (5×10^6^) with 5×10^6^ autologous (n=3) or allogeneic (n=11) PBMCs. Allogeneic PBMCs were either whole (n=4), depleted of CD8^+^ cells (n= 3) or depleted of CD56^+^ cells (n=4). Cell depletions were performed prior to injection using a magnetic bead-based technique. 7 days following injection, cells were recovered from the peritoneal cavity by lavage. CD4^+^CFSE^+^ Tregs in the effluent were enumerated by flow cytometry. Cell numbers are plotted for each mouse individually, with mean ± SD for each treatment group. (**D**) The absolute number of Tregs. Statistical significance was calculated using t-tests: autologous versus allogeneic PBMCs (p=0.0001), whole versus CD8^+^-depleted PBMCs (p=0.0008) and whole versus CD56^+^-depleted allogeneic PBMCs (p=0.8188). (**E**) The absolute number of undivided Tregs. Statistical significance was calculated using t-tests: autologous versus allogeneic PBMCs (p=0.0002), whole versus CD8^+^-depleted PBMCs (p=0.0008) and whole versus CD56^+^-depleted allogeneic PBMCs (p=0.2179). (**F**) The percentage of CFSE^hi^ (representing undivided) Tregs. Statistical significance was calculated using t-tests: autologous versus allogeneic PBMCs (p=0.0342); whole versus CD8^+^-depleted PBMCs (p=0.0011) and whole versus CD56^+^-depleted allogeneic PBMCs (p=0.0983)

To confirm the role of host alloimmunity on Treg survival, we investigated the effect of HLA-matching of Tregs with the host. We screened >150 Treg/PBMC pairs to identify matches between Class I HLA (-A, -B, -C) and Class II HLA (-DP, -DQ, -DR). Three donor pairs were evaluated: B218-B209, B150-B208, and B130-B209 with HLA mismatches of (0,1,1,2), (0,0,0,1), and (1,2,1,2), respectively, across the 6 examined loci. This provided a range of partially matched Treg-recipient pairs **(Table S1)**. Skin allograft experiments were performed as before but mice were treated with a reduced Treg dose to create a more challenging model (*25*). Mice were treated with either partially matched or partially/completely mismatched allogeneic Tregs **(Fig. 3A)**. While mice receiving completely and partially mismatched allogeneic Treg promptly rejected their grafts with an MST of 24 days and 27 days respectively **(Fig. 3B, C)**, most mice receiving partially matched allogeneic Treg developed long-term graft survival (>100 days). Two animals in the partially mismatched group developed xenoGvHD and were censored from the analysis from day 33, further illustrating the reduced efficacy of partially mismatched Tregs.

**Fig. 3.**
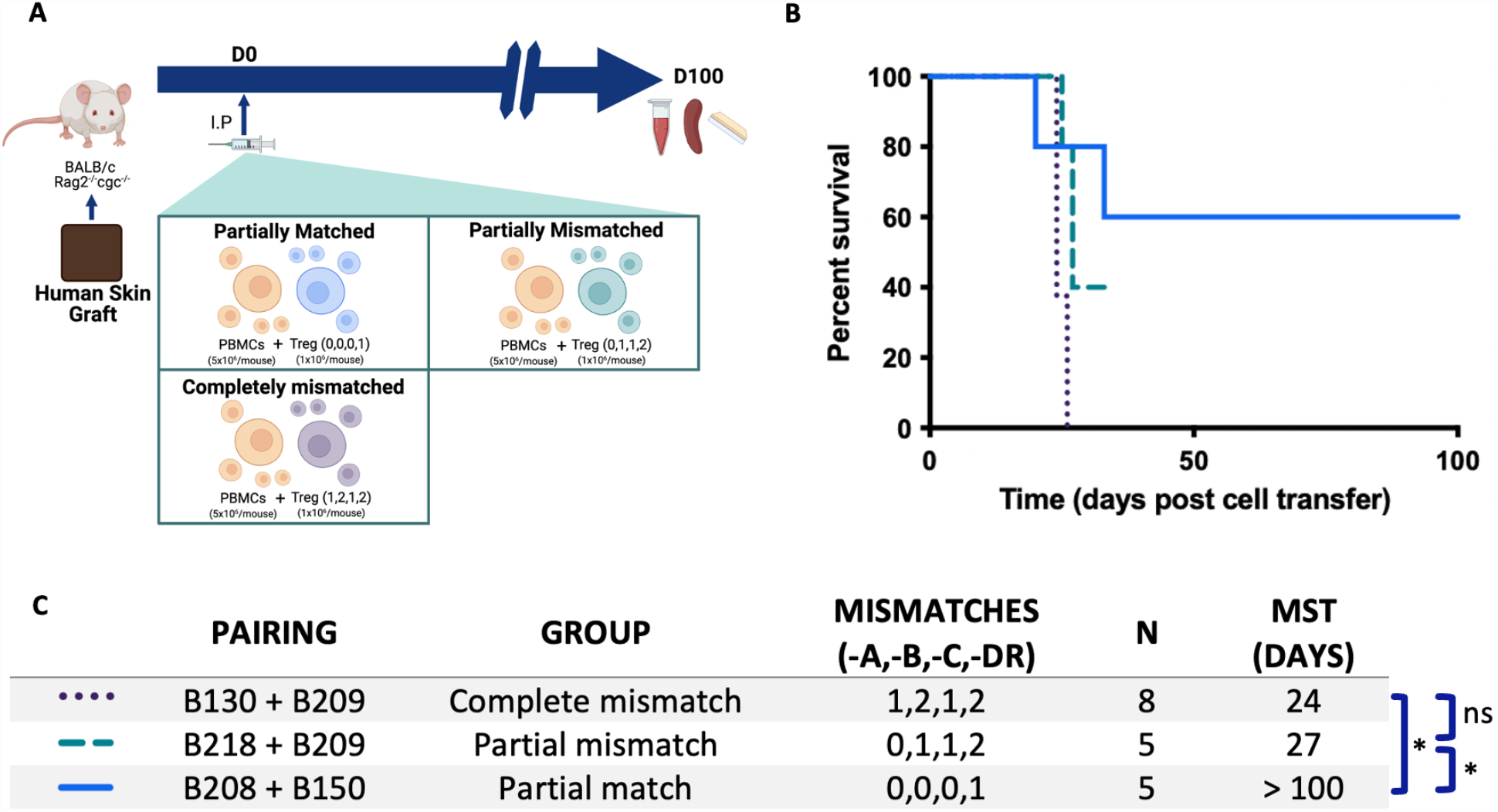
HLA matching of adoptively transferred allogeneic Tregs to the host enables long-term human skin transplant survival. (**A**) Immunodeficient BALB/c Rag2^-/-^ cγc^-/-^ mice grafted with human skin received intraperitoneal human PBMCs (5×10^6^) and either completely mismatched (1,2,1,2, n=8), partially mismatched (0,1,1,2, n=5), or partially matched (0,0,0,1, n=5) Tregs (1 × 10^6^) at HLA-A,-B,-C,-DR loci. Grafts were monitored for macroscopic signs of rejection over the subsequent 100 days. (**B**) Percentage of grafts surviving is plotted over time post-adoptive transfer of cells. The mice receiving partially mismatched Treg developed signs of xenogeneic graft-versus-host-disease necessitating removal of these animals from the experiment at day 33. **(C)** Tabulated median survival time (defined as the time at which half of the grafts were rejected). Groups were analyzed using the Mantel-Cox log-rank test with multiple testing correction using the Benjamini-Hochberg method (all groups χ^2^=7.84, df=2, p=0.02; complete mismatch versus partial mismatch χ^2^=6.02, df=1, p=0.01; complete mismatch versus partial match χ^2^=4.37, df=1, p=0.0364; partial mismatch versus partial match χ^2^=0.33, df=1, p=0.57)

### Generation of hypoimmunogenic Treg by gene editing of key genes for HLA-class I and II

Identification of HLA-matched donors for patients requires complex logistics and complete matching remains a challenge (*26, 27*). Therefore, we explored whether targeted genetic modification of HLA may alleviate the need for stringent HLA-matching (**Fig. 4**). CRISPR-Cas9-mediated genetic disruption of *beta-2-microglobulin* (*B2M*) alone (II, *B2M* KO) and combined with deletion of *class II, major histocompatibility complex, transactivator* (*CIITA*) (III, double KO) eliminated the expression of HLA-class I and class II, respectively (**Fig. 4A-D**). As *B2M*-edited T cells were shown to be targeted by NK cells via missing-self activation (*28*), we established a gene editing strategy to introduce the NK-cell inhibitory receptor HLA-E into Tregs. By integrating the coding sequence of the non-polymorphic *HLA-E* gene with a short linker sequence into the *B2M* exon 2, we installed an HLA-E-*B2M* fusion gene in HLA-class I-negative Treg (IV, HLA-E KI) (**Fig. 4B-E**). To further eliminate HLA-class II in *B2M*-edited HLA-E knock-in (KI) Treg, we silenced *CIITA* using a CRISPR-Cas9-derived adenine base editor (ABE) (*29*) in a second manipulation step (V, HLA-E KI/CIITA KO, H/C, Treg) (**Fig. 4B-E)**. HLA-engineered Tregs displayed the canonical features of Treg identity including a CD4^+^CD25^+^FOXP3^+^ phenotype (**Fig. 4F**), and no elevated Th1 cytokine production after polyclonal stimulation (**Fig. 4G)**. All gene-edited Treg products suppressed the proliferation of allogeneic T cells in a dose-dependent manner and were comparable to autologous control Tregs in an *in vitro* suppression assay (**Fig. 4 H,I**). Gene editing did not alter the low methylation state at the Treg-specific demethylation region (TSDR, **Fig. 4J**) in *FOXP3*.

**Fig. 4.**
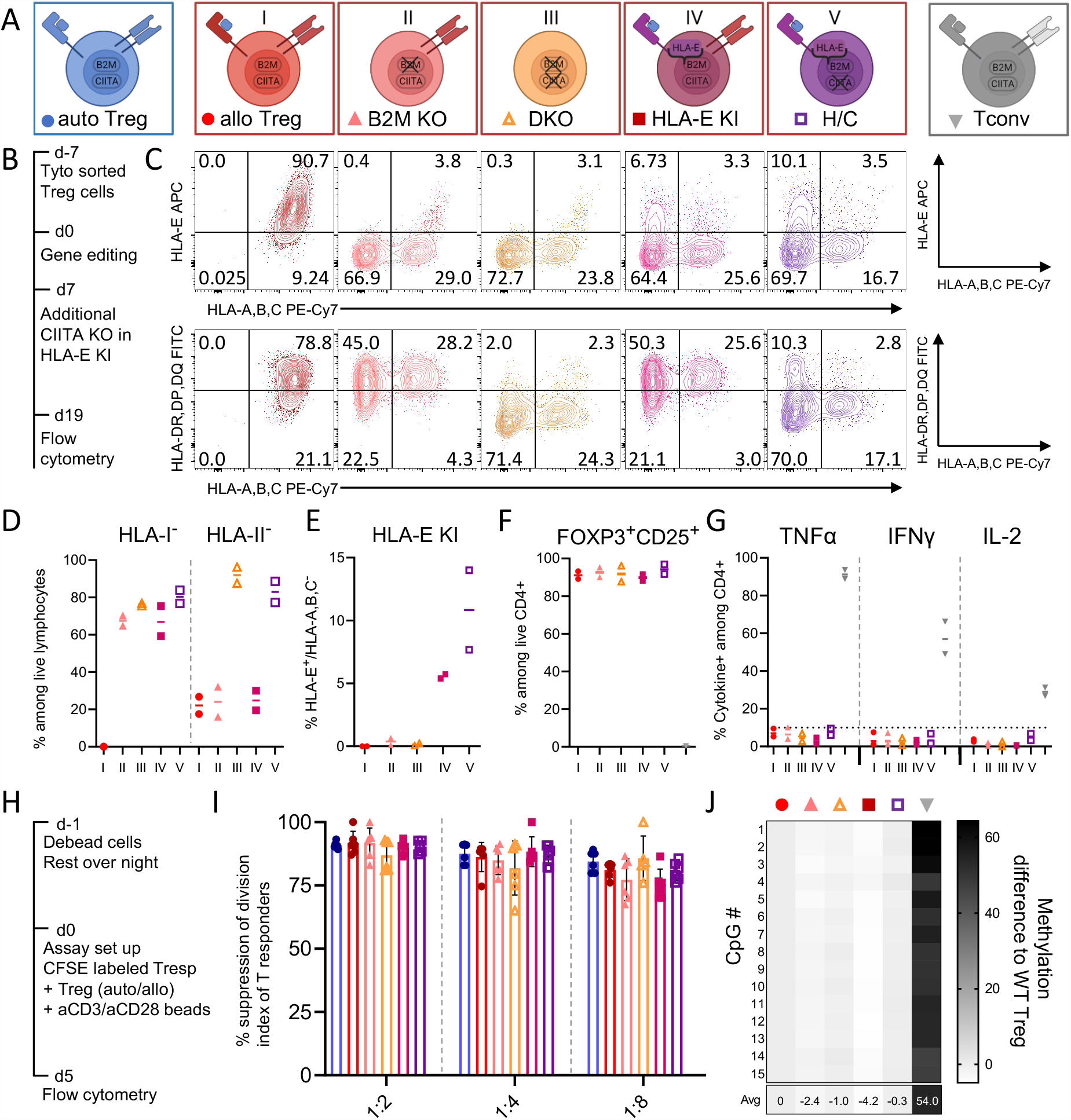
HLA gene-edited allogeneic Tregs maintain their canonical identities *in vitro*. (**A**) Gene editing strategy using CRISPR/Cas9 to disrupt HLA class-I and class II by targeting *B2M* and/or *CIITA*, respectively. HLA-E was inserted with a short linker sequence into *B2M* exon 2, generating an HLA-E-*B2M* fusion protein. For H/C Tregs the additional *CIITA* KO was performed by splice site disruption using the adenine base editor ABE-8.20-m. (**B**) Timeline of cell isolation, gene-editing and surface protein expression measurement. (**C**) Representative contour plots of editing outcome by HLA-A,B,C (PE-Cy7), HLA-DR,DP,DQ (FITC) and HLA-E (APC) staining. (**D**) Summary of editing outcome for two biological replicates showing frequency of HLA class-I and class-II negative cells and (**E**) of HLA-E positive and HLA-A,B,C negative knock-in cells among live lymphocytes. (**F**) Treg identity was confirmed by staining of CD25 and intracellular staining of FoxP3 as well as (**G**) intracellular cytokine (TNFα, IFNγ, IL-2) expression after PMA/Ionomycin stimulation with conventional T cells as controls. (**H**) Timeline of suppression assay: CSFE-stained human CD3^+^ cells (2.5×10^4^) with and without either autologous or the different allogeneic Treg conditions (at 1:1, 1:4 and 1:8 Treg:Tresp ratios) were stimulated with αCD3αCD28-coated beads at a 1:1 ratio of beads to total cells (Treg+Tresp). (**I**) CSFE dilution after 5 days incubation was measured by flow cytometry and a division index was calculated. (**J**) Targeted DNA-methylation analysis of the 15 CpGs within the *FOXP3-*TSDR using bisulfite amplicon sequencing. Data were normalized by subtracting the methylation frequency of the WT Treg sample. Mean of two biological replicates is shown.

### Transgenic HLA-E partially protects from NK cell-mediated lysis in vitro

Fusing the non-polymorphic *HLA-E* cDNA with the endogenous *B2M* gene prevented surface expression of polymorphic HLA-class I complexes in Tregs (**Fig. 4B**), but it should also reduce missing-self activation and cytolysis by endogenous NK cells (*30*). To test this hypothesis, we generated primary NK cell lines from healthy human donors and expanded them *in vitro* using cytokine-containing medium (**Suppl. Fig. S3A**,**B)**. When co-culturing the pre-activated NK cells with Tregs, we observed an NK cell significant dose-dependent cytolysis of Tregs with deleterious edits at the *B2M* locus (**Suppl. Fig. S3C**). Analysis of the remaining Tregs confirmed dose-dependent lysis of HLA-class I-negative Tregs (**Suppl. Fig. S3D-F**), but a preferential survival of HLA-class I-positive and HLA-class I-negative, HLA-E-*B2M* KI Treg (**Suppl. Fig. S3E-G**). The relative fold-increase of HLA-class I-positive Tregs was slightly higher than the relative enrichment observed for HLA-E, indicating only partial protection from NK cell killing.

### Manipulation of HLA-class I and II maintains effectiveness of allogeneic Tregs in vivo

To test whether our HLA-engineered Treg cells are protected from allospecific T cell rejection, we established allospecific T cell lines by stimulating allogeneic CD56-depleted PBMCs with irradiated T cell-depleted PBMCs from our Treg donors as targets (**Suppl. Fig. S4A**). Prior to re-stimulation with allogeneic target cells, the alloreactive T cells were enriched for CD3^+^ cells to remove contaminating NK cells and other CD3^-^ cells (**Suppl. Fig. S4B**). The allospecific T cells comprised >90% TCRα/β^+^ T cells with CD4^+^ and CD8^+^ T cell subsets and no NK cells (**Suppl. Fig. S4B**,**C**). Allospecific T cells induced dose-dependent lysis of the unmodified allogeneic Tregs (**Suppl. Fig. S4G**,**H**). In 3/5 allospecific T cell lines, we observed similar lysis of unmodified as well as allogeneic Tregs which were only edited in the *B2M* locus (II-*B2M* KO, IV-HLA-E-*B2M* KI only) (**Suppl. Fig. S4H**). Tregs with silenced HLA-class I and II were significantly better protected from allospecific T cell lysis than Treg edited at *B2M* alone (**Suppl. Fig. S4I**). These data suggest that allospecific CD4^+^ T cells can contribute to the rejection of unmatched allogeneic Tregs in an HLA-class II-dependent manner.

Finally, we tested HLA-engineered Tregs *in vivo* under challenging conditions (**Fig. 5A**) with a complete mismatch (0/10 HLA-match) between donors for humanization and the Treg donor as well as the lower 1:5 Treg:PBMC ratio. To mimic a true ‘off-the-shelf’ scenario, the Treg products were cryopreserved after manufacturing and thawed immediately prior to injection as previously described (*31*). As expected, allogeneic Tregs were unable to prevent the rapid rejection of transplanted human skin grafts (**Fig. 5B**). Fully edited allogeneic Tregs with HLA-E-*B2M* KI and *CIITA* KO (H/C Treg) promoted graft survival in 40% of the animals, comparable to autologous control Tregs (**Fig. 5B)**. Our results emphasize the necessity to silence or match both HLA Class I and II to protect allogeneic Treg products from mismatched donors.

**Fig. 5.**
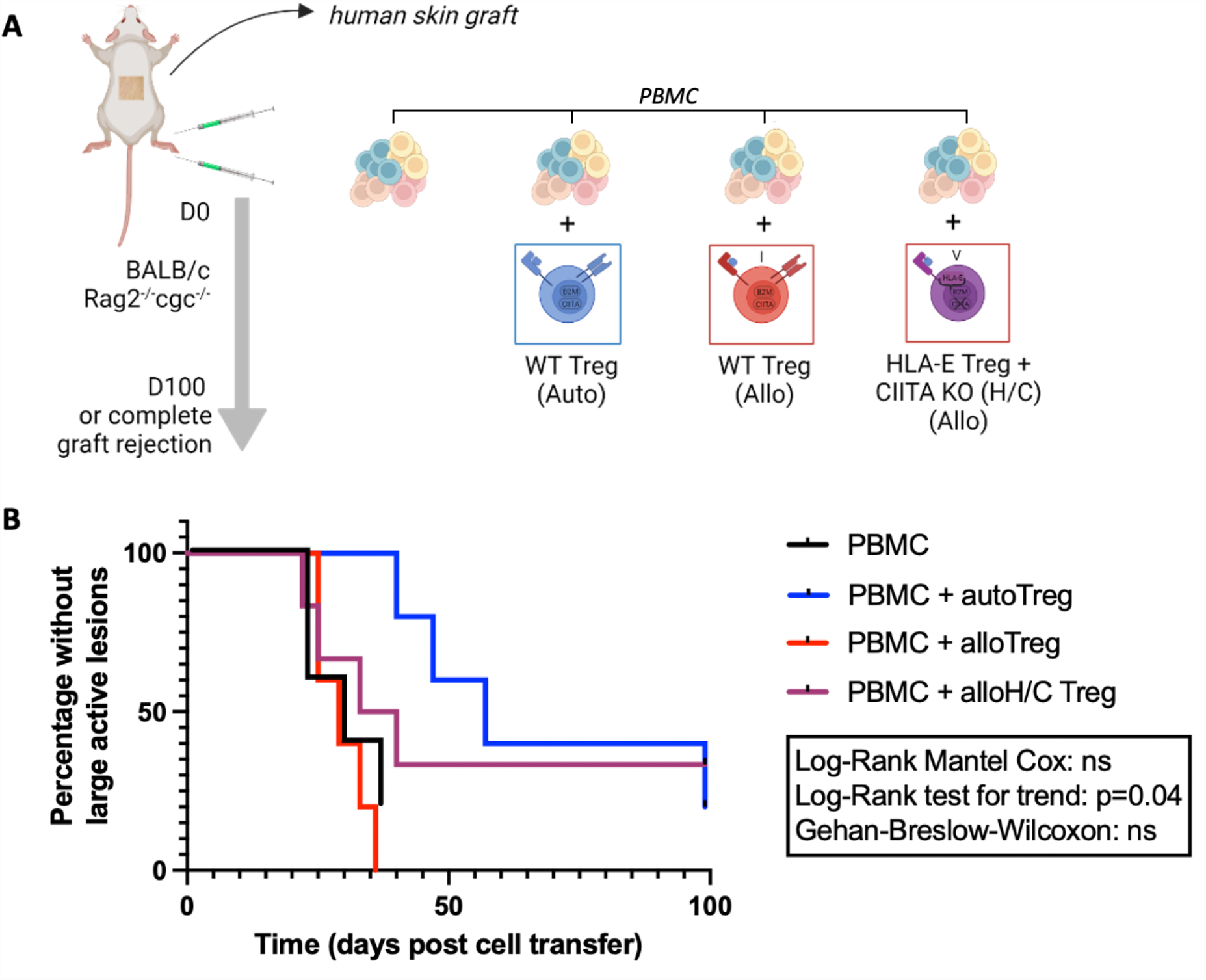
HLA-engineered Tregs without polymorphic HLA-I and -II enable long-term protection of human skin transplants from rejection, comparable to autologous Tregs. (A) Immunodeficient BALB/c Rag2^-/-^ cγc^-/-^ mice grafted with human skin received intraperitoneal human PBMCs (5×10^6^, n=5) alone, with Tregs autologous to the PBMC donor (1×10^6^ Treg, n=5), with Treg allogeneic to the PBMC donor (1×10^6^ Treg, n=5), or with allogeneic HLA class 2 KO and HLA-E KI Treg (1×10^6^, n=5). Grafts were monitored for macroscopic signs of rejection over the following 100 days. (F) Percentage of grafts without large active lesions is plotted over time post-adoptive transfer of cells. Statistical significance was assessed using Mantel-Cox log rank tests.

## Discussion

Mounting evidence demonstrates the therapeutic efficacy of autologous Treg therapy in a variety of pathologies mandating consideration of how this cell therapy approach may be scaled to deliver maximal benefit to all patient populations. To date, most clinical trials have been conducted with small numbers of patients in academic-sponsored efforts with manufacture of cell products by small GMP-compliant in-house units (*3*). For example, manufacture of the autologous Treg product for our on-going Phase 2b trial involves numerous manual steps undertaken by highly trained scientists (*9, 32*); simply put, to increase the number of products produced in this facility would require a linear increase in scientists, facilities, and equipment. From an economic perspective, therefore, there is currently limited potential to reduce costs by scaling up production. This may significantly hamper transition of the technology into the pharmaceutical industry and increase the risk of traditional funding institutions rejecting the approach in its entirety. Beyond the technical elements, the unpredictability inherent in producing a biological ‘living’ product introduces significant variability in both the input and output of what is otherwise a standardized manufacturing process. Moreover, Treg cell therapy production is carried out over several weeks, making rapid treatment with an autologous product impossible for newly diagnosed conditions (e.g. new onset diabetes), or where the timing of treatment cannot be planned (e.g. deceased donor organ transplantation). Several clinical trials of regulatory cell therapy have been prematurely terminated due to manufacturing difficulties with the cellular product (*33*).

Here, we have demonstrated that Tregs suppress in an HLA-agnostic fashion, with equal potency towards autologous and allogeneic effector cells *in vitro*. Unlike conventional effector T cells, allogeneic Tregs do not result in the development of GvHD *in vivo* (*34, 35*). As expected, allogeneic Tregs were subject to CD8^+^-mediated depletion *in vivo*, which effectively reduced the number of cells available to exert an immunosuppressive effect. Depletion of CD8^+^ T cells but not CD56^+^ NK cells restored survival and the immunoregulatory potency of unmatched allogeneic Tregs *in vivo*. By creating an immunologically inert environment at the time of Treg administration, either by stringent HLA-matching (9/10) or gene editing of HLA class I and II, we demonstrate that Tregs survive for long enough to establish a regulatory environment, conferring a similar degree of allograft tolerance to autologous cells. This provides the first evidence that the therapeutic efficacy of allogeneic Tregs can be significantly enhanced by matching and/or gene editing paving the way for an improved ‘off-the-shelf’ strategy to mass manufacture an allogeneic Treg cellular therapy.

These findings are congruent with the current paradigm of Treg suppressive mechanisms which are largely TCR/HLA-independent, including expression of co-inhibitory molecules such as cytotoxic T lymphocyte antigen (CTLA-4), production of immunomodulatory cytokines such as TGF-β, IL-10, IL-35, preferential utilization of IL-2 through expression of the high-affinity IL-2 receptor, and induction of apoptosis in effector populations through granzyme expression (*1*). Both upstream expression of inhibitory ligands or the production of inhibitory cytokines, and their respective downstream receptors, are HLA-independent and therefore *ex vivo* Treg activation alone would be expected to promote an immunosuppressed milieu. Further, the Treg population contains cells with alloreactive T cell receptors (*36*) which could boost the immunomodulatory function, because such alloreactive Tregs were shown to be preferentially activated in the allograft and draining lymph nodes in mice models (*37, 38*). Despite the consistent *in vitro* data, we also highlight the challenges with the use of *in vitro* suppression assays to predict *in vivo* function. Here, we observe equal suppressive potency between autologous and allogeneic Tregs *in vitro*, supporting the notion that cell-intrinsic mechanisms enable allogenic Treg to exert a suppressive effect irrespective of their donor. However consistent performance of these cells in vitro did not necessarily translate to efficacy *in vivo*.

The development of hypoimmunogenic, gene-edited ‘off-the-shelf’ cellular therapies has been proposed for tissue replacement (*23, 39*), anti-tumor cell therapy with conventional T cells (*24, 40-42*), and now, allogeneic Treg applications. Silencing of HLA on allogeneic cells may avoid allosensitization and allow redosing of allogeneic cell products in case of relapse or for consolidating treatment (*43*). Introduction of NK-cell inhibitory receptors will be required to improve the persistence of allogeneic Treg with disrupted HLA-class I expression (*43*). We demonstrate that an HLA-E-*B2M* fusion gene allows partial protection against NK cells. HLA-E is a non-classical HLA class Ib molecule with only two alleles present in diverse populations (HLA-E*01:01 and HLA-E*01:03) (*44*). HLA-E primarily presents signal peptides from other HLA-I molecules; however, some reports also demonstrate that HLA-E*01:03 can present viral peptides from CMV or EBV (*45, 46*). Binding of HLA-E to CD94/NKG2A or CD94/NKG2B on NK cells or CD8 T cells inhibits said cells, explaining the protective effect observed *in vitro* (*47*). In contrast, HLA-E can also present viral peptides to activate NK cells by binding the conserved NKG2C (*48*) or EBV-specific and HLA-E-restricted CD8^+^ T cells (*46, 49*). Previous strategies to create hypoimmunogenic cells implemented HLA-E fusion proteins that lack antigen-presentation by covalently linking B2M, HLA-E and a non-polymorphic signal peptide of HLA-G into a single molecule (*23, 24*). Our HLA-E-*B2M* fusion gene is based on HLA-E*01:03 without a blocking peptide and therefore may retain some ability to present pathogenic peptides. However, due to the severely compromised antigen presentation, the first clinical application of HLA-engineered hypoimmunogenic Tregs cell product may warrant inclusion of a genetic safety switch.

Allogeneic Treg therapy requires installation of multiple edits to reduce immunogenicity of HLA-class I and II in a universal cell source, such as iPSCs, and in *ex vivo* expanded Tregs from healthy human donors. The latter requires effective gene editing at multiple sites. Targeting multiple genes with conventional CRISPR-Cas9 provokes high rates of translocations with unknown biological function (*50, 51*). To overcome this, we separated the editing step between KI of HLA-E and silencing of *CIITA* to eliminate HLA-II with an ABE enzyme. Base editors, such as the employed ABE (*29*), introduce targeted base changes without inducing DNA breaks and thereby reduce the risk of gross genetic rearrangements in the gene edited cell product. In future, our editing strategy may be adapted to enable multiplex-editing in a single manipulation to reduce manufacturing using different nucleases (*52*) or novel gene editing tools for DSB-free KI (*53, 54*).

Allogeneic Tregs offer a promising therapeutic approach, but their immunosuppressive capacity is hindered by the anticipated alloresponse. Here we pave the way for two viable strategies to enable their effective use. The first approach involves the use of HLA-matched allogeneic Tregs, while the second strategy incorporates unmatched Tregs with HLA class I and class II KO, alongside the introduction of HLA-E. Our results demonstrate that both of these approaches facilitate the long-term survival of Tregs *in vivo* resulting in a therapeutic effect.

This study therefore presents a significant advancement towards the clinical application of ‘off-the-shelf’ Treg cell therapy derived from universal donors. Not only do our findings provide a path to overcoming the current hurdles in allogeneic Treg therapy, but they also open the door to realizing the full potential of this transformative therapeutic approach for a variety of immune-mediated pathologies.

## Funding

European Union, Horizon 2020, grant agreement: 825392 (FI, JH, OM, WD, VG, JKP, MSH, HDV, PR, DLW)

ERC starting grant ‘EpiTune’, grant agreement 803992 (JKP)

Medical Research Council, Clinical Research Training Fellowship, award reference: MR/V000942/1 (OM)

Wellcome Trust 211122/Z/18 (FI)

Kidney Research UK (JH)

British Heart Foundation, FS/ 12/72/29754 (KM)

Berlin Institute of Health, SPARK-BIH (JK, DLW)

## Author contributions

Conceptualization: OM, WD, VG, JH, FI, DLW

Data curation: OM, KM, WD, VG, CF, MY, JKP, JH, FI, DLW

Formal Analysis: OM, WD, VG, CF, MY,

Funding acquisition: JH, FI, JKP, HDV, PR, DLW

Investigation: OM, KM, WD, VG, CF, JK, MV, MY

Methodology: OM, KM, WD, VG, CF, JK, MV, AK, JKP, MSH, PR, HDV, DLW

Project administration: JH, FI, MSH, HDV, PR, DLW

Resources: AK, JKP, MSH, PR, HDV, JH, FI, DLW

Software: not applicable Supervision: JH, FI, DLW

Validation: OM, WD, VG, CF, JK, MV

Vizualisation: OM, WD, VG, JH, FI, DLW

Writing—original draft: OM, WD, VG, HDV, JH, FI, DLW

Writing—review & editing: OM, WD, VG, HDV, JKP, JH, FI, DLW

## Data and materials availability

All data are available in the main text or the supplementary materials.

