## Supplementary Methods and Materials for "Matching or genetic engineering of HLA Class I and II facilitates successful allogeneic ‘off-the-shelf’ regulatory T cell therapy"

### Materials and Methods

Cell isolation and cell culture

Human peripheral blood mononuclear cells (PBMC) were isolated from the blood of healthy donors (provided by the National Blood Service, Oxford, U.K) or was obtained after informed and written consent (Charité ethics committee approval EA4/091/19) using Lymphosep (Biowest, Nuaillé, France) for density gradient centrifugation. Erythrocytes were lysed using PharmLyse buffer (BD biosciences, Franklin Lakes, NJ, USA). Pre-enrichment of $CD25^+$ cells from PBMCs was performed using CD25 microbeads (Miltenyi Biotech, Bergisch Gladbach, Germany) according to the manufacturer's protocol. Depletion of $CD8^+$ cells or $CD56^+$ cells from PBMCs was performed using αCD8 or αCD56 microbeads (Miltenyi Biotech) according to the manufacturer's protocol. Treg ($CD4^+CD25^+CD127^{lo}$) were FACS-sorted from CD25-enriched PBMCs using BD FACSAria (BD Biosciences) after staining with αCD4 – ECD (Beckmann Coulter, Brea, CA, USA), αCD25 – PE-Cy7 (BD Biosciences), and αCD127 – PE (BD Biosciences). $CD4^+$ effector T cells (Teffs) were isolated from CD25-depleted PBMCs using αCD4 microbeads (Miltenyi) according to the manufacturer's protocol. Cells were cryopreserved in freezing medium comprising 50% heat-inactivated FCS and 50% RPMI1640 with 10% DMSO. Prior to assay cells were thawed in a 37°C water bath before dilution of DMSO with RPMI1640 warmed to 37°C.

For experiments intended for gene-editing, pre-enrichment of $CD4^+$ cells from PBMCs was performed using CD4 microbeads (Miltenyi Biotech) according to the manufacturer´s protocol. $CCR7^+$ Tregs ($CD4^+CD25^+CD127^{low}CCR7^+$) were sorted using a Tyto sorter (Miltenyi Biotech) after staining with αCD4 VioBlue (Miltenyi, REAL103), αCD25 APC (Miltenyi, REAL128), αCD127 PE-vio770 (Miltenyi, REAL102), αCD45RA FITC (Miltenyi, REAL164) and CCR7 PE (Biolegend, G043H7). Sorted Tregs were cultured in a 96 U well plate with 100,000 cells in 200µl Treg medium per well. Treg medium consists of X-Vivo 50 (Lonza) medium supplemented with 10% heat-inactivated FCS, 500IU/mL of recombinant human interleukin-2 (IL-2) (Miltenyi, Bergisch Gladbach, Germany) and 100nM Rapamycin (Pfizer). T cells were cultured in T cell medium (RPMI1640 supplemented with 10% FCS, 10ng/mL IL-7, and 5ng/mL IL-15). NK cells were enriched from PBMCs using the NK isolation Kit (Miltenyi) and cultured in NK MACS Medium (Miltenyi) supplemented with 10% FCS, IL-2 (500IU /mL) and IL-15 (5ng/mL). The K562 cell line was cultured in RPMI 1640 supplemented with 10% FCS.

For *in vitro* cell cultures and assays, leukocytes were cultured in RPMI1640 supplemented with L-glutamine (Sigma-Aldrich, St. Louis, MO, USA), 100U/mL penicillin and 10mg/mL streptomycin (Sigma-Aldrich), and 10% heat-inactivated human serum (Seralab, Haywards Heath, U.K).

Treg expansion

After isolation, Tregs were cultured for 16 days in complete medium supplemented with 1000IU/mL of recombinant human IL-2 (Novartis Pharmaceuticals UK Ltd, Surrey, UK). Treg were stimulated with αCD3αCD28 T cell activator beads (ThermoFisher Scientific, Waltham, Massachusetts, USA) on day 0 (at a ratio of three beads to one cell) and day 7 (at a ratio of one bead to one cell). Cultures were passaged and medium changed as required. Cells were rested without beads in medium supplemented with of 200U/mL IL-2 for forty-eight hours prior to assay.

For experiments intended for gene-editing, isolated Tregs were initially stimulated with MACS® GMP ExpAct™ Treg Kit at a bead:cell ratio of 4:1 (Miltenyi Biotech, Bergisch Gladbach, Germany) one day after isolation. Five days following isolation, cells were re-stimulated at a 1:1 bead:cell ratio. Beads were removed prior to non-viral gene editing using a strong magnet stand. Electroporated cells were rested overnight without beads before re-stimulation at a 1:1 bead:cell ratio. For further expansion rounds, Tregs were split and re-stimulated on alternate days at a 1:1 bead:cell ratio. All gene edited and unedited Tregs were cryopreserved on day 23 following cell isolation and were thawed as required.

Generation of HLA-E knock-in construct

A dsDNA homology directed repair template (HDRT) was used for the targeted insertion of HLA-E into the *B2M* locus, creating a B2M-HLA-E fusion protein and disrupting the endogenous *B2M* gene. Homology arms mediating the insertion into B2M exon 2 flanked the HDRT. The construct contained the remaining part of B2M exon 2 as well as exon 3 followed by a flexible (G4S)4 linker and the HLA-E transgene (Table S2). HDRTs were generated as previously described (*1*). In brief, multiple fragment InFusion cloning was performed according to the manufacturer's protocol (Clontech, Takara) with purified PCR fragments (Kapa Hotstart HiFi Polymerase Readymix, Roche) and using synthesized DNA (gBlocks, IDT). In-Fusion cloning strategies were planned with SnapGene (Insightful Science; snapgene.com). Sequence validation of HDR-donor-template-containing plasmids was performed by Sanger Sequencing (LGC Genomics, Berlin). The B2M-HLA-E template was amplified from the plasmid by PCR using the primers (B2M_F: 5′ aagctcatttggccagagtgg 3′ and B2M_R: 5′ agctagaggaagccagtaggtaag 3′). PCR products were purified and concentrated using paramagnetic beads (AMPure XP, Beckman Coulter Genomics). HDRT concentrations were quantified using the Qubit 4 fluorometer (Thermo Fisher Scientific) and a Qubit™ dsDNA BR-Assay-Kit according to the manufacturer's protocol and adjusted to 1µg/mL in nuclease-free water.

Gene editing to modulate HLA surface expression on primary human Treg

After 7 days of culture MACS® GMP ExpAct™ Treg Kit beads were depleted using a MACSiMAG™ Separator (Miltenyi). Cells were removed from the magnet, counted, and then washed twice in sterile PBS by centrifugation at 100×g for 10 min at room temperature (RT). In parallel, Cas9 RNP was prepared. For the electroporation of $10^6$ primary Tregs, 0.5µL of poly(L-glutamic acid) (PGA) (molecular weight 15,000-50,000, Sigma-Aldrich, 100µg/µL), 0.48µL of synthetic modified sgRNA (Table S3; 100µM in TE buffer; IDT), and 0.4µL recombinant SpCas9 protein (Alt-R S.p. Cas9 Nuclease V3; IDT; 61µM) were mixed by thorough pipetting. The mixture was incubated for 15 min at RT and placed on ice. For HLA-E KI conditions, 0.5µL of HDRT (stock concentration: 1µg/µL) was added prior to electroporation. $10^6$ harvested Tregs were resuspended in 20µL ice-cold P3 electroporation buffer (Lonza) just before electroporation to keep the exposure time to the electroporation buffers to a minimum. 20µL of resuspended cells were transferred to the RNP/HDRT suspension, mixed thoroughly, and transferred into a 16-well electroporation strip (Lonza) without any air bubbles. The cells were electroporated using the EH-115 program on the 4D-Nucleofector (Lonza). Immediately after electroporation, 90µL of pre-warmed Treg medium was added per well. After 10 min, the cells were carefully resuspended and transferred to two 96-well round-bottom plates (50µL/well) containing 150µL pre-warmed Treg medium per well. The gene editing of HLA-E KI into *B2M* plus KO of *CIITA* Tregs (H/C Tregs) was performed in two subsequent editing steps. 7 days after HLA-E KI into *B2M* a second electroporation was performed using adenine base editor ABE8.20-m (*2*) mRNA produced in house by in vitro transcription as described previously (*3*). Tregs were harvested, beads were magnetically removed, and cells were washed twice in PBS. $5x10^6$ cells were resuspended in

100µl P3 electroporation buffer (Lonza) and mixed with 2µg of ABE8.20-m mRNA and 0.48µL of synthetic modified sgRNA (100µM in TE buffer; IDT). The suspension was electroporated in 100µl Nucleocuvette Vessels (Lonza) using a Lonza 4D nucleofector device (program EH-115). 900µL of pre-warmed Treg medium was added per cuvette. After 10 min, the cells were carefully resuspended and transferred to a 24 well cell culture plate.

Flow Cytometry

7-AAD viability staining solution (Affymetrix, Santa Clara, CA, U.S.) or fixable blue dead cell stain kit (Thermo Fisher, U.S.) were used to eliminate dead cells from analysis. For phenotyping the following antibodies (supplier, clone) were used: mouse anti-human CD127 PE (BD, hIL-7R-M21), mouse anti-human CD25 PE-Cy7 (BD, M-A251), mouse anti-human CD3 eFluor450 (Affymetrix, OKT3), mouse anti-human CD4 ECD (Beckmann Coulter, SFCI12T4D11), mouse anti-human CD45 APC (BD, RPA-T4), mouse anti-human CD45 APC (Affymetrix, H130), mouse anti-human CD8 FITC (Affymetrix, SK1), mouse anti-human CD8 PE (BD, HIT8a), mouse anti-human CD8 APC-Cy7 (BD, SK1), rat anti-human FOXP3 FITC (Affymetrix, PCH101), rat anti-human FOXP3 PE (Affymetrix, PCH101), rat anti-human FOXP3 eFluor450 (Affymetrix, PCH101), rat anti-mouse CD45 PE (Affymetrix, 30F11), mouse anti-human HLA-A,B,C PE-Cy7 (Biolegend, W6/32), mouse anti-human HLA-E APC (Biolegend, 3D12), mouse anti-human HLA-DR,DP,DQ FITC (Biolegend, Tü39), mouse anti-human FOXP3 FITC (BD Pharmingen, 259D/C7), mouse anti-human CD25 PC7 (Beckman, B1.49.9), mouse anti-human TNFα A700 (Biolegend, Mab11), mouse anti-human IFNγ APC-eF780 (Invitrogen, 4S.B3), rat IL-2 PE-Cy7 (Biolegend, MQ1-17H12), mouse anti-human CD4 PE (Beckman, 13B8.2), mouse anti-human CD8 PE-Cy7 (BD Pharmingen, RPA-T8), mouse anti-human CD3 PB (Biolegend, UCHT1), mouse anti-human CD56 A647 (Biolegend, 5.1H11), recombinant human anti-human NKG2A FITC (Miltenyi, REA110), recombinant human anti-human NKG2C PE (Miltenyi, REA205). Fluorescence was measured using either BD FACSCanto or Beckman Coulter CytoFLEX cytometers.

*in vitro* suppression assay

Responder PBMCs were stained with 10µM CFSE (Thermo Fisher Scientific, Waltham, MA, USA) according to the manufacturer's protocol and cultured ($1x10^5$ per well) for 72hrs with autologous or allogeneic *in vitro*-expanded Treg, at ratios ranging from 1:1 to 1:8 Treg:responders. αCD3αCD28 T cell activator beads (Thermo Fisher Scientific) were added at a 1:5 bead:cell ratio. All assays were performed in triplicate. For analysis of responder proliferation from CFSE dilution profiles, division indices were calculated as previously described (*4*). The number of Treg ($7AAD^-CFSE^-CD3^+CD4^+$ cells) surviving following assay was expressed as a proportion of the initially plated cells. Treg enumeration was performed for 6 assays each using Treg from separate donors with 2-3 replicates per condition.

Mice

BALB/cRag2$^{-/-}$cγc$^{-/-}$ mice (Jackson Laboratory, Bar Harbor, ME, USA) were housed in individually ventilated cages in the John Radcliffe Hospital Biomedical Services Unit under specific pathogen-free conditions. All protocols were conducted in accordance with the UK Animals (Scientific Procedures) Act (1986) and approved by Oxford University's Committee on Animal Care and Ethical Review.

Humanized Mouse Model of Skin Allograft Rejection

Human skin was procured with full informed written consent and with ethical approval from the Oxfordshire Research Ethics Committee (REC B), study number 07/H0605/130. Surgical grafting of a 1cm$^2$ human split-thickness skin graft onto the flank of BALB/cRag2$^{-/-}$cγc$^{-/-}$ recipients was performed as previously described (*5*). A total of 68 age and sex matched mice were used in two allograft survival experiments. Five weeks following skin grafting, mice received 5x10$^6$ cryopreserved PBMC in pure RPMI via intra-peritoneal injection, with or without 1x10$^6$ or 5x10$^6$ *in vitro*-expanded Treg. Treg were either autologous or allogeneic to the PBMC donor. Both PBMCs and Treg were allogeneic to the skin graft. Grafts were observed for macroscopic markers of rejection by an assessor blinded to treatment group. Mice were sacrificed following either graft rejection or at day 100 post-transplantation (whichever came sooner) by cervical dislocation. Blood was harvested from the inferior vena cava following schedule one for flow cytometric analysis. Treatments were allocated evenly between cages and litters. Raw data processing was performed by a researcher blinded to treatment group.

<u>*in vivo* mixed leukocyte assay</u>

Treg were stained with 10µM CFSE (Thermo Fisher Scientific). BALB/cRag2$^{-/-}$cγc$^{-/-}$ mice received 5x10$^6$ cryopreserved PBMCs in pure RPMI via intraperitoneal injection with or without 5x10$^6$ *in vitro*-expanded CFSE-labelled Treg. For each experiment, Treg were isolated from a single donor with allogeneic PBMC isolated from a second. Flow cytometric counting beads (Thermo Fisher Scientific) were co-injected with cells (5x10$^5$ per mouse) to enable normalisation of absolute cell counts between lavage samples. 7 days after injection of cells, peritoneal lavages were performed by flushing the peritoneal cavity with 10mL saline following midline laparotomy and collecting the effluent. Cells extracted by lavage were analysed by flow cytometry. For enumeration and phenotypic analyses, Treg were identified in lavage fluid by flow cytometry and CFSE staining. A total of 44 mice were used for mixed leucocyte assays, all of whom were female and aged between 8-12 weeks.

<u>HLA-matching</u>

Peripheral blood underwent full tissue typing (HLA-A, -B, -C, -DP, -DR, and -DQ) by the Oxford Transplant Centre Histocompatibility and Genetics Laboratory.

<u>TSDR analysis</u>

FOXP3-TSDR methylation analysis was performed by bisulfite amplicon sequencing as previously described (*6*). Briefly, genomic DNA was isolated using the Quick-DNA Microprep Kit (Zymo Research D3020, Irvine, USA) according to the manufacturer´s protocol. Up to 200 ng genomic DNA was bisulfite-converted using EZ-DNA methylation Gold kit (Zymo Research D5005, Irvine, USA). Subsequently PCRs were performed (10 µl of bisulfite-treated DNA, 2xKAPA HiFi Hotstart Uracil+ ReadyMix (Kapa Biosystems, USA, KK2802), 0.25 mM of each dNTP, using 0.3 pmol of primers (F1: 5' ACACTCTTTCCCTACACGACGCTCTTCCGATCTTTTGGGGGTAGAGGATTTAGAG GG-3' and R3: 5'GACTGGAGTTCAGACGTGTGCTCTTCCGATCTCCACATCCACCAACACCCAT - 3'). Amplicons were purified with QIAquick PCR Purification Kit (Qiagen Germany, 28106), normalized to 20ng/ul, and sequenced (2x300 bp paired-end). Reads were aligned and evaluated using the bismark package (*7*) ( https://pubmed.ncbi.nlm.nih.gov/21493656/).

<u>Generation of alloreactive T cell lines</u>

PBMCs were isolated from three healthy donors. CD56$^+$ cells were depleted from whole PBMCs using CD56 microbeads (Miltenyi) according to the manufacturer's protocol. As feeder cells, CD3-depleted PBMCs from two Treg donors were irradiated (30Gy). Feeder cells were cultured with CD56-depleted PBMCs at a ratio of 1:1 for 9 days in T cell medium (RPMI1640 supplemented with 10% FCS, 10ng/ml IL-7 and 5ng/mL IL-15). CD3 enrichment was performed on day 9 prior to a second round of stimulation with irradiated feeder cells at a ratio of 1:1 until day 20. T cells were split every other day. Lineage phenotyping was performed using αCD3 Pacific Blue, αCD56 AF647, αβTCR PE, γδTCR FITC, αCD4 APC-Fire750 and αCD8 BV510 (see above for details). Alloreactive T cells were cultured in cytokine-free medium overnight prior to the cytotoxic assay.

*in vitro* alloreactive T cell cytotoxic assay

Expanded alloreactive T cells were co-cultured with wild type or gene-edited allogeneic Tregs. Alloreactive T (AlloT) cells were stained with CFSE (Thermo Fisher Scientific) prior to co-culture. AlloT cells were added to 25,000 target T cells in 96-well, round-bottom, cell culture plates at various alloT:Treg ratios (2:1, 4:1, 8:1) with control wells containing only the target cells. Plates were centrifuged at 100g for 1 min at RT and incubated at 37°C and 5% CO2. A 96 well cell culture plate containing 50µL of PBS with DAPI (final dilution 1:100,000, stock concentration: 1 mg/mL, Thermo Scientific) was prepared and cooled to 4°C for 30 min prior to stopping the assay. 22h after setting up the co-culture, 50µL of resuspended cells were added to the 50µL of PBS/DAPI solution followed by a minimum of 10 minutes incubation at 4°C. 30µL of the cell suspension were analyzed by flow cytometry to evaluate the number of viable cells. Alloreactive T cell-mediated cytotoxicity was calculated as the reduction in cell number of CFSE-negative target cells in the co-culture normalised to the target only control.

*in vitro* primary NK cell cytolytic assay

NK cells were freshly isolated from the PBMCs of four healthy donors using the NK isolation kit (Miltenyi, Gemany) according to the manufacturer's instructions. Isolated NK cells were cultured in NK medium and were expanded prior to phenotypic characterization by flow cytometry (see above for info). Six days following isolation, expanded NK cells were used in the *in vitro* cytotoxicity assay. In brief, 100,000 target cells (gene edited Treg, unedited Treg, or K562 controls) were co-cultured at various NK:target cell ratios (0,25:1, 0,5:1, 1:1) for 20h before flow cytometry. Of the 200µL cell suspension, 50µL was used to establish the number of viable target cells whilst 150µL was used to establish target HLA-A,B,C (PE-Cy7) and HLA-E (APC) expression.

Statistical Analysis

All data were analysed, and graphs produced, using GraphPad Prism version 9.0 for MacOS (GraphPad Software, La Jolla California USA). For *in vitro* Treg suppression assays repeated measures two-way ANOVAs were used to assess statistical significance, with the Bonferroni *post hoc* correction applied for pairwise comparisons between treatments. Student's T-test or Mann-Whitney U test were applied to comparisons of cell frequencies between autologous and allogenic PBMC/Treg combinations. Graft survival kinetics were assessed using Mantel-Cox log rank tests for each pairwise comparison between groups. All plotted error bars represent mean ± standard error of the mean (SEM) unless otherwise stated.

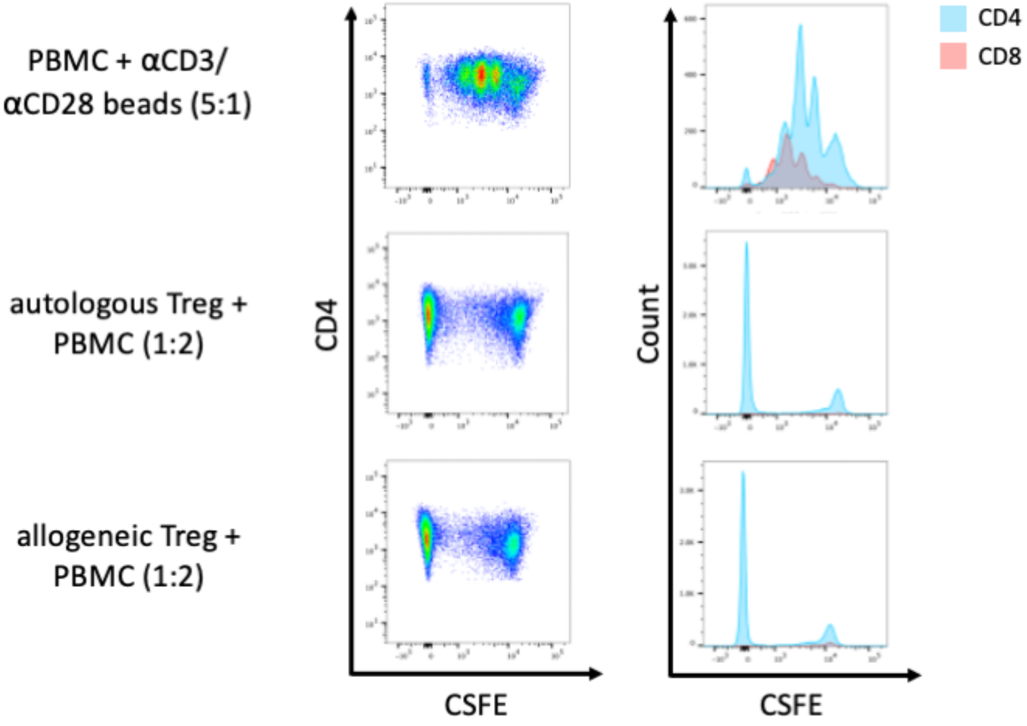


**Fig. S1. Representative flow cytometry plots of *in vitro* suppression of T Cell proliferation and activation by autologous versus allogeneic Tregs**

Responder PBMCs were stained with CFSE and cultured at $1x10^5$ per well for 72h with autologous or allogeneic *in vitro*-expanded Treg. αCD3αCD28 T cell activator beads (Thermo Fisher Scientific) were added at a 1:5 bead:cell ratio. Representative flow cytometry dot plots of $CD4^+$ CSFE signal dilution (left column) and histograms of $CD4^+$ and $CD8^+$ CSFE intensity (right columnn) for each condition.

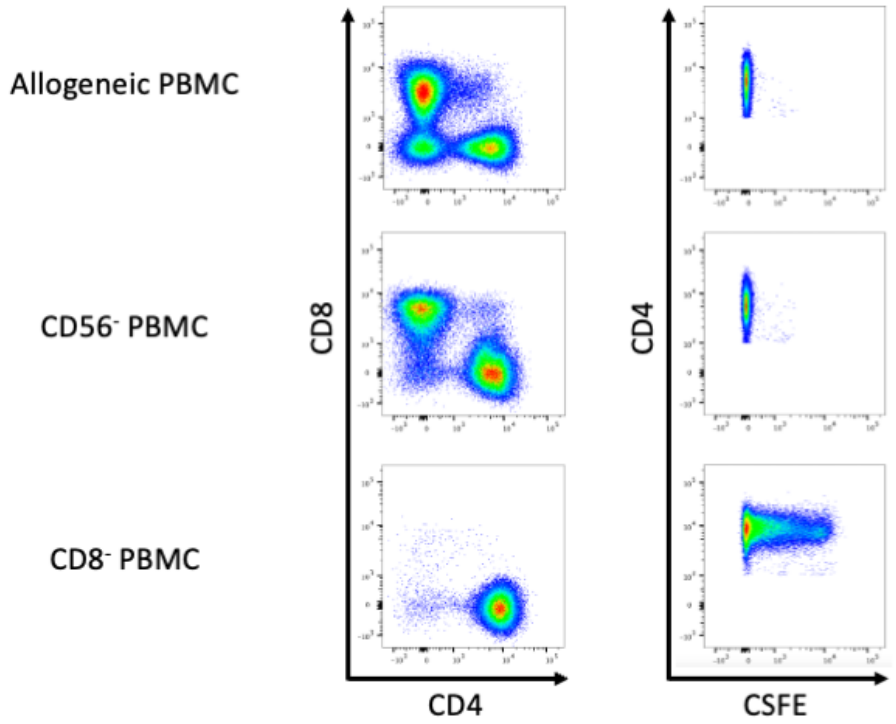


**Fig. S2. Representative flow cytometry plots of *in vitro* suppression of CD56- or CD8-depleted PBMC by third-party Treg**

BALB/cRag2$^{-/-}$cγc$^{-/-}$ mice received 5x10$^{6}$ CSFE-stained Treg with 5x10$^{6}$ of either whole PBMC, CD8-depleted PBMC, or CD56-depleted PBMC in pure RPMI via intraperitoneal injection. 7 days after injection, cells were recovered from the peritoneal cavity by lavage. Human lymphocytes in the effluent were enumerated and phenotyped by flow cytometry. Representative plots of CD4$^{+}$ and CD8$^{+}$ staining within the hCD45$^{+}$CD3$^{+}$ live lymphocyte gate (left column) and CSFE staining within hCD45$^{+}$CD3$^{+}$CD4$^{+}$ live lymphocyte gate (right column, representing Treg).

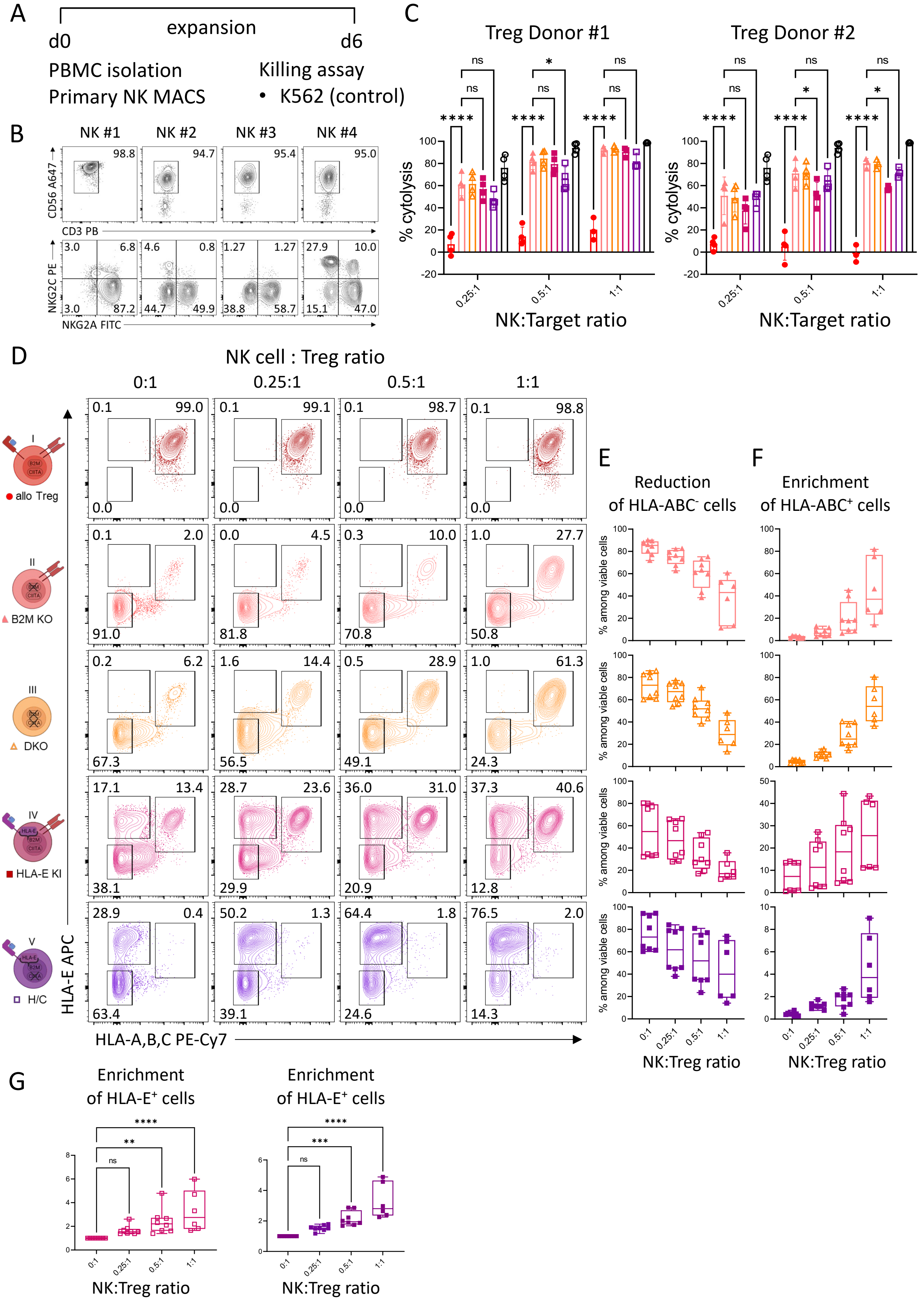


**Fig. S3. Co-culture of primary expanded NK cell lines with gene edited Treg demonstrates partial evasion of NK mediated missing-self cytolysis by expression of transgenic HLA-E.**

(**A**) NK cells were MACS isolated and expanded for 6 days. (**B**) Phenotype of the NK cells isolated from peripheral blood of 4 healthy donors by flow cytometric analysis of CD56 AF647, CD3 PE, NKG2C PE and NKG2A FITC. (**C**) Cytolysis of Tregs and K562 cells by NK cells normalized to target only control after co-culture at NK:target cell ratios of 0.25:1, 0.5:1 and 1:1. (**D**) Representative contour plots of HLA-A,B,C and HLA-E staining upon 20 hours co-culture of primary NK cells and Treg cells at various ratios: 0:1, 0.25:1, 0.5:1, 1:1. NK cells were pre-labelled with CFSE. Red represents allo Treg cells; pink presents B2M KO Treg cells; golden presents DKO Treg cells; magenta represents HLA-E KI Tregs; and purple represents HLA-E KI and CIITA KO (H/C) Tregs. (**B**) Percentage of HLA-A, B, C negative cells among viable cells at various ratios of NK cells to Tregs. N=6-8. (**C**) Percentage of HLA-A, B, C positive cells among viable cells at various ratios of NK cells to Tregs. N=6-8. (**D**) Enrichment of HLA-E positive cells when co-culturing of Tregs with NK cells at various ratios. (N=6-8). For statistic analyses ordinary two-way ANOVAs with main effects only were used to compare each column mean with the control column (C: B2M KO; G: NK:Treg ratio 0:1). Dunnett testing was performed for statistical hypothesis testing (ns: $p \geq 0.05$; *: $p < 0.05$; **: $p < 0.01$; ***: $p < 0.001$; ****: $p < 0.0001$).

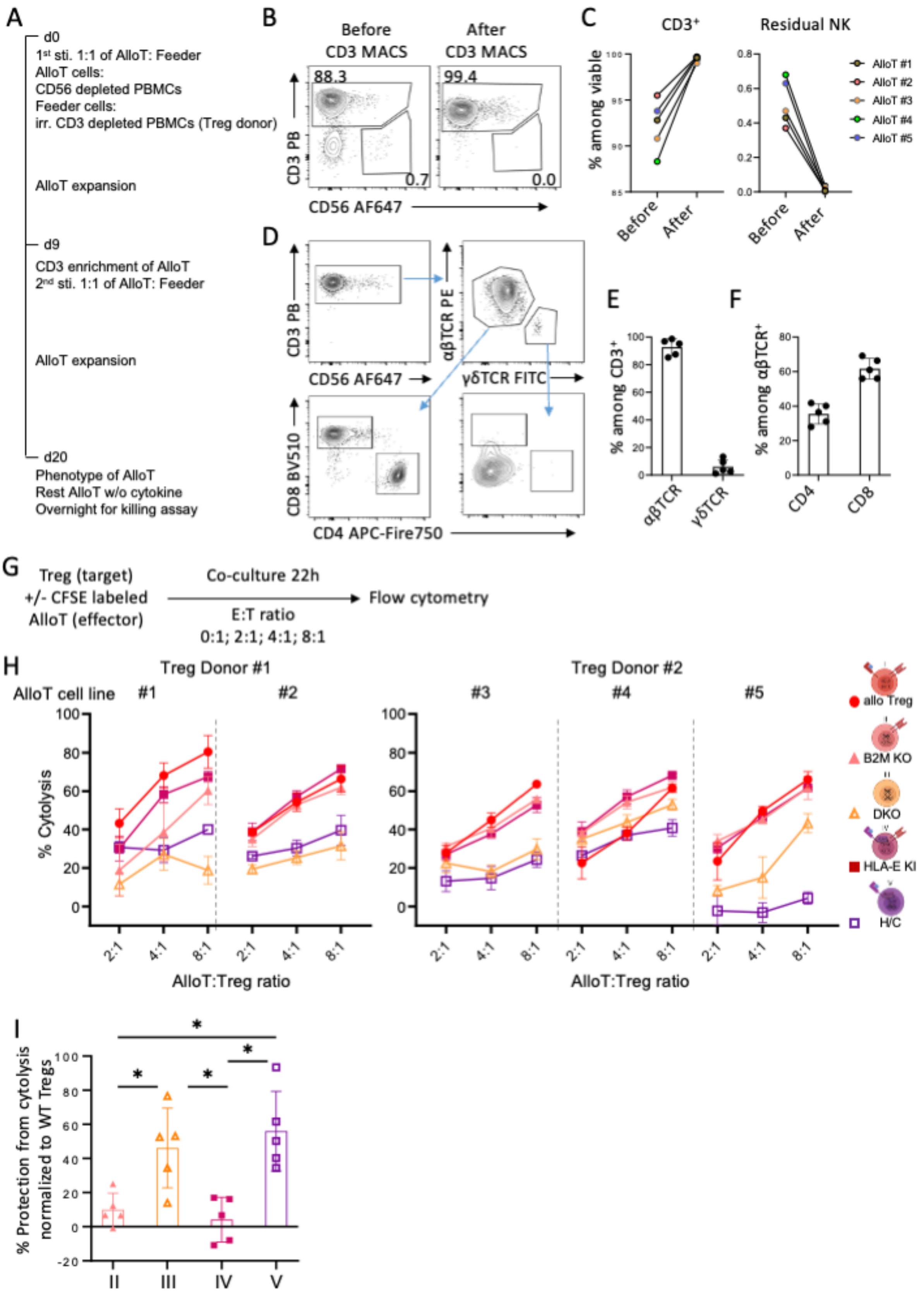
A
d0
1st sti. 1:1 of AlloT: Feeder
AlloT cells:
CD56 depleted PBMCs
Feeder cells:
irr. CD3 depleted PBMCs (Treg donor)
AlloT expansion
d9
CD3 enrichment of AlloT
2nd sti. 1:1 of AlloT: Feeder
AlloT expansion
d20
Phenotype of AlloT
Rest AlloT w/o cytokine
Overnight for killing assay
B
Before CD3 MACS
After CD3 MACS
CD3 PB
CD56 AF647
C
CD3+
Residual NK
% among viable
Before
After
AlloT #1
AlloT #2
AlloT #3
AlloT #4
AlloT #5
D
CD3 PB
CD56 AF647
αβTCR PE
γδTCR FITC
CD8 BV510
CD4 APC-Fire750
E
% among CD3+
αβTCR
γδTCR
F
% among αβTCR+
CD4
CD8
G
Treg (target)
+/- CFSE labeled
AlloT (effector)
Co-culture 22h
E:T ratio
0:1; 2:1; 4:1; 8:1
Flow cytometry
H
Treg Donor #1
Treg Donor #2
AlloT cell line
% Cytolysis
AlloT:Treg ratio
allo Treg
B2M KO
DKO
HLA-E KI
H/C
I
% Protection from cytolysis normalized to WT Tregs
II
III
IV
V

**Fig. S4. KO of HLA-class I and class II and HLA-E KI protects Tregs from allogeneic T cell cytolysis.**

**(A)** Allogeneic T cells were generated over three weeks with two rounds of stimulation by irradiated (30Gy) CD3-depleted PBMCs from the Treg donor prior to killing assay. **(B)** Representative contour plots of CD56 AF647 and CD3 PB staining of the allogenic T cells before and after CD3 MACS enrichment at day 9 prior to the second round of stimulation. **(C)** Percentages of $CD3^+$ T cells (left) and of residual $CD56^+CD3^-$ NK cells (right) of five allogenic T cells before and after CD3 MACS enrichment (n=5). **(D-F)** Allogenic T cell phenotype one day prior to killing assay. **(D)** Representative contour plots of CD3 PB, CD56 AF647, αβTCR PE, γδTCR FITC, CD4 APC-Fire750, CD8 BV510. **(E)** Percentages of αβTCR and γδTCR amongst $CD3^+$ T cells (n=5). **(F)** Percentages of $CD4^+$ and $CD8^+$ lymphocytes amongst $αβTCR^+$ T cells (n=5). **(G)** Schematic overview of the allogenic T cell killing assay. Allogeneic T cells (effector, E) were pre-labelled with CFSE prior to co-culturing with Tregs (target, T) at E:T ratios of 0:1, 0.25:1, 0.5:1, 1:1 for 22 hours. **(H)** Cytolysis of allogeneic Treg cells with different gene editing strategies from two donors by allo-specific T cell lines. Each dot represents mean of three technical replicates. Percentage cytolysis was normalized to cell counts from Treg cells alone (without allo-specific T cells) = E:T ratio 0:1. (I) Relative protection of different gene edited Treg products in comparison to unedited allogeneic Treg at 8:1 E:T ratio combining all data from the five experiments shown in H. For statistical analysis one-way ANOVA with Geisser-Greenhouse correction was performed. To adjust for multiple corrections, follow up test was performed with the Tukey test (ns: $p \geq 0.05$; *: $p < 0.05$ **: $p < 0.01$). Only statistically significant results are indicated.

| Alleles | PBMC | | Tregs | | Mismatches |
|---|---|---|---|---|---|
| | **B208** | | **B150** | | |
| A | 2 | 1 | 2 | 1 | **0** |
| B | 62 | 8 | 62 | 8 | **0** |
| C | 9 | 7 | 9 | 7 | **0** |
| DR | 17 | 11 | 13 | 17 | **1** |
| DR | 52 | | 52 | | |
| DQ | 2 | 7 | 2 | 6 | |
| | **B218** | | **B209** | | |
| A | 3 | 2 | 3 | 2 | **0** |
| B | 44 | 7 | 35 | 7 | **1** |
| C | 5 | 7 | 4 | 7 | **1** |
| DR | 4 | 12 | 1 | 15 | **2** |
| DR | 52 | 53 | 51 | | |
| DQ | 7 | | 5 | 6 | |
| | **B130** | | **B209** | | |
| A | 2 | 29 | 3 | 2 | **1** |
| B | 8 | 44 | 35 | 7 | **2** |
| C | 16 | 7 | 4 | 7 | **1** |
| DR | 17 | 7 | 1 | 15 | **2** |
| DR | 52 | 53 | 51 | | |
| DQ | 2 | | 5 | 6 | |

**Table S1. HLA typing of PBMC and Treg donor pairs**

PBMCs and Tregs were HLA-typed by the Oxford Transplant Centre Histocompatibility and Genetics Laboratory. HLA loci for all experimental pairings are presented for each HLA allele. For each allele, the number of mismatches are quantified. The B130/B209 pair is completely mismatched at HLA-A,-B,-C,-DR (1,2,1,2); B218/B209 is partially mismatched at these loci (0,1,1,2); and B208/B150 is partially matched (0,0,0,1).

**Table S2.**

Homology-Directed DNA Repair Template (HDRT) used for HLA-E knock-in into *B2M locus*

**Table S3.**

CRISPR-Cas9 guide RNAs used for editing of *B2M* and *CIITA*
